# Reverse remodelling of the mitochondria and cytoskeleton after respiratory heart rate variability pacing of the failing sheep heart

**DOI:** 10.1101/2025.05.29.656916

**Authors:** D J Crossman, G Guo, J Shanks, M Pachen, J Bai, H Moammer, M Middleditch, A C Grey, J Paton, R Ramachandra

## Abstract

We have previously demonstrated that pacing the failing sheep heart with respiratory heart rate variability (RespHRV), a natural variability in the heart rate that is linked to respiration on a breath-by-breath basis, improves cardiac output dramatically. In this study, we used proteomics and super-resolution microscopy to explore the role of energetics and T-tubule cellular remodelling in response to ResPHRV pacing in an ischaemic ovine model of heart failure (HF). After 2 weeks of RespHRV pacing, cardiac output improved by 1.1 ± 0.2 L/min (**p=0.003). Sequential Window Acquisition of all Theoretical Mass Spectra (SWATH-MS) was used to probe differences between three groups: HF without any intervention, HF with RespHRV pacing and a healthy control group. Orthogonal Partial Least Squares (OPLS) discriminant analysis demonstrated a clear separation of all three groups by T score (***p<0.001) with the HF+RespHRV pacing group placed intermediate between the HF and control groups. The top 50 proteins negatively correlated with T score (down in HF, restored after RespHRV) were dominated by mitochondrial proteins, as confirmed by Pathway Enrichment Analysis (***p<0.001). Multiple Reaction Monitoring Mass Spectrometry (MRM-MS) analysis confirmed this finding in selected targets (ACAA2, ACADS, CRAT, NDUFA8, and SUCLG1, *p<0.05). STimulated Emission Depletion (STED) microscopy identified a disruption of mitochondria structure in HF (*p<0.05) that was restored in the HF+R group (p=0.051). The area of mitochondria labelling was increased in the HF+RespHRV group compared to HF (**p=0.005). Many cytoskeletal proteins linked to mitochondria regulation and T-tubule remodelling were upregulated in HF and were reduced by RespHRV. MRM-MS was able to confirm these findings for selected targets (ANAXA2, CAVIN2, SPTBN1, TUBA4A). STED microscopy of collagen VI and the ryanodine receptor revealed cellular hypertrophy and remodelling of the T-tubules and cardiac junctions in HF sheep (*p<0.05), RespHRV showed a trend for reversing these structural changes. These data support the hypothesis that within the first two weeks of RespHRV pacing, there is an increase in mitochondrial repair and function coupled with re-organisation of the cellular cytoskeleton, which is consistent with the improvement in cardiac pump function.

## Introduction

In all species, the heart has evolved to beat at non-regular intervals, yet pacemakers pace metronomically. A variable heart rate is a positive indicator of health and is (in most part) linked to respiration^1^, a phenomenon originally called respiratory sinus arrhythmia^2^ but herein named respiratory heart rate variability (RespHRV). RespHRV is most prominent in the young and physically fit and lost in cardiovascular disease. We have developed a novel cardiac pacemaker that restores RespHRV every breath. Our pre-clinical studies demonstrate RespHRV pacing increases cardiac output in both a rodent and an ovine HF model^3^ ^4^. In sheep with HF induced by microembolisation, RespHRV pacing drives a dramatic 22% increase in cardiac output (1.5 L/min) that peaks after ∼7 days of pacing, is sustained over 4 weeks of pacing and once pacing was switched off, declines slowly over 7 days^3^. The improvement in cardiac output is not due to a hemodynamic change as responses to RespHRV pacing were not immediate but appeared 4 days after pacing onset, suggesting a genomic response. How RespHRV improves cardiac function is not understood. However, our previous mathematical model indicated that RespHRV may enhance cardiac function through improved energetics^5^. The heart is a highly oxidative organ and dysfunctional energetics, particularly decreased mitochondrial respiration, is a major contributor to cardiovascular disease, with ATP levels reduced by ∼30% in HF^6^. Thus, we hypothesized that the restoration of cardiac energetics is a potential mechanism behind the increase in cardiac output with RespHRV pacing.

In our previous study in HF sheep, RespHRV pacing reduced myocyte hypertrophy and reverse remodelled the transverse tubules (T-tubules)^3^, a cellular structure critical for Ca^2+^ cycling and contractility. Cardiac hypertrophy is a known risk factor for poor outcomes in human heart failure^7^. Moreover, we have previously demonstrated that disrupted T-tubule structure is highly correlated with the loss of contractile function within the failing human heart^8^, suggesting improvement in these two features could, in part, underpin the improved cardiac output after RespHRV pacing. Both pathological cardiac hypertrophy and loss of T-tubules reduce energetic efficiency in rat cardiac muscle^9,10^ providing a potential link between our modelling work^5^ and the cellular improvements we have documented after RespHRV pacing^3^. To gain further insight into the cellular and molecular pathways involved in the response to RespHRV pacing, we undertook a proteomic and super-resolution microscopy study of the left ventricle (LV) in HF sheep after 2-weeks of RespHRV pacing and compared to the LV of HF sheep without pacing as well as a cohort of healthy sheep. Our central hypothesis is that RespHRV improves the energetics of the heart by restoring metabolic pathways and improving the organisation of the T-tubular and mitochondrial networks.

## Methods

### Animal model

Adult (3–5 year old) female Romney sheep were housed in individual crates, and acclimatized to laboratory conditions (18°C, 50% relative humidity, 12 h light–dark cycle) and human contact for 1 week before any experiments. Sheep were fed 2–2.5 kg/day (Country harvest pellets) and had access to water ad libitum. All animal experiments and surgical procedures followed relevant guidelines and were approved by the Animal Ethics Committee of the University of Auckland (#2082). Experiments were carried out over a period of 6 months in each animal.

Three groups of sheep were studied; normal healthy (C), non-paced HF (HF), and HF paced with RespHRV (HF+R). Animals were randomly allocated to the RespHRV or the no intervention groups. HF failure was induced as previously described^11,12^ and is detailed below. Echocardiography was performed (GE Healthcare Vivid™ S70N Ultrasound) to measure ejection fraction in conscious healthy animals (typically 70–80%). To induce heart failure with reduced ejection fraction, sheep were anaesthetized with 2% Diprivan (Propofol) (5 mg/kg IV AstraZeneca, AUS), maintained with a 2% isoflurane-air-O_2_ mixture and were intubated for mechanical ventilation. The left or right femoral artery was accessed percutaneously using an 8F (CORDIS®, USA) sheath and under fluoroscopic guidance, the left main coronary artery cannulated, and the catheter advanced into either the proximal left main coronary artery or left descending coronary arteries. The animals in the two HF groups underwent sequential weekly (1–3 weeks) embolizations using polystyrene latex microspheres (45 μm; 1.2 mL, approximately 650 microspheres, Polysciences, Warrington, PA, USA). Prior to the injection of microspheres, β-blocker (metoprolol up to 20 mg/kg, IV) and lignocaine (2 mg/kg,IV) were injected intravenously to prevent ventricular arrhythmias. An electrocardiogram (ECG) was recorded from lead II prior to the infusion of the microspheres and for a further 5 min after infusion. The recordings were made on a dual bio amp electrocardiograph switch box with a power lab and LabChart (AD Instruments, NZ). A change in the ST segment (elevation or depression) and T wave (inversion) was taken to indicate a successful embolization. Three days post embolization, conscious sheep underwent echocardiography; once the ejection fraction dropped to ∼ 45%, no more embolizations were performed.

RespHRV pacing and recording of cardiac output were performed as previously described^12^. In brief, three months after heart failure was induced by microembolization sheep underwent instrumentation under general anaesthesia (induced with 2% Diprivan (Propofol) 5 mg/kg IV, AstraZeneca, AUS) and maintained with a 2% isoflurane-air-O_2_ mixture. An intercostal nerve block was performed on ribs 2–6, using a 22G needle 1 mL sterile saline followed by 3 mL Bupivavaine which was injected into the intercostal space immediately cranially of each rib. A Doppler flow probe (Size 22, Transonic, AU) was placed around the ascending aorta for a direct measure of cardiac output on a beat-by-beat basis. For cardiac pacing, two pacing leads (Biotonik, Berlin, Germany, Solia S 53-in case of failure in one) were secured externally to the left atrium with 3.0 Filasilk suture and silicone gel. To gain a real-time measure of respiration, electrodes were implanted into the diaphragm to give a measure of diaphragmatic EMG. Sheep were given antibiotic injections (6 mL IM; Oxytetra, Phenix, NZ), and analgesia (Ketofen 10%, 1 mL IM; Merial, Boehringer Ingelheim, NZ) at the start of surgery, and for the first 3 days post-operatively. Animals were allowed to recover for 5–7 days before start of the protocol.

Cardiovascular and respiratory parameters were recorded simultaneously from conscious sheep with heart failure on a desktop computer with a CED micro 1401 interface and a data acquisition program (Spike2 v8, Cambridge Electronic Design, UK). A baseline recording period was acquired when heart rate and cardiac output had stabilized post-operatively (between 5 and 7 days). Continuous arterial blood pressure, cardiac output from the ascending aorta blood flow probe and heart rate were recorded 24 h per day. Heart rate was calculated from the inter-pulse interval of the blood flow in the ascending aorta. DEMG signal was amplified (X10,000), and filtered (band pass 0.3–3.0 kHz). RespHRV pacing was achieved using a biofeedback device described previously^4,12,13^. This device performed real-time integration of the diaphragmatic electromyographic (EMG) activity^14^, used to define the inspiratory phase (Ceryx Medical, UK; Fig. 1A). Pacing was set 10–15 beats per minute above resting heart rate with an RSA magnitude (peak-to-trough) of 12 beats per minute. Pacing voltage was set at 1.5 V and pulse width at 2 ms. Pacing voltage was increased if and as needed during the two-week pacing period. RespHRV pacing was visually checked against the DEMG channel to ensure the rising phase of heart rate correlated with inspiration, and the falling phase of heart rate with expiration.

**Figure 1.**
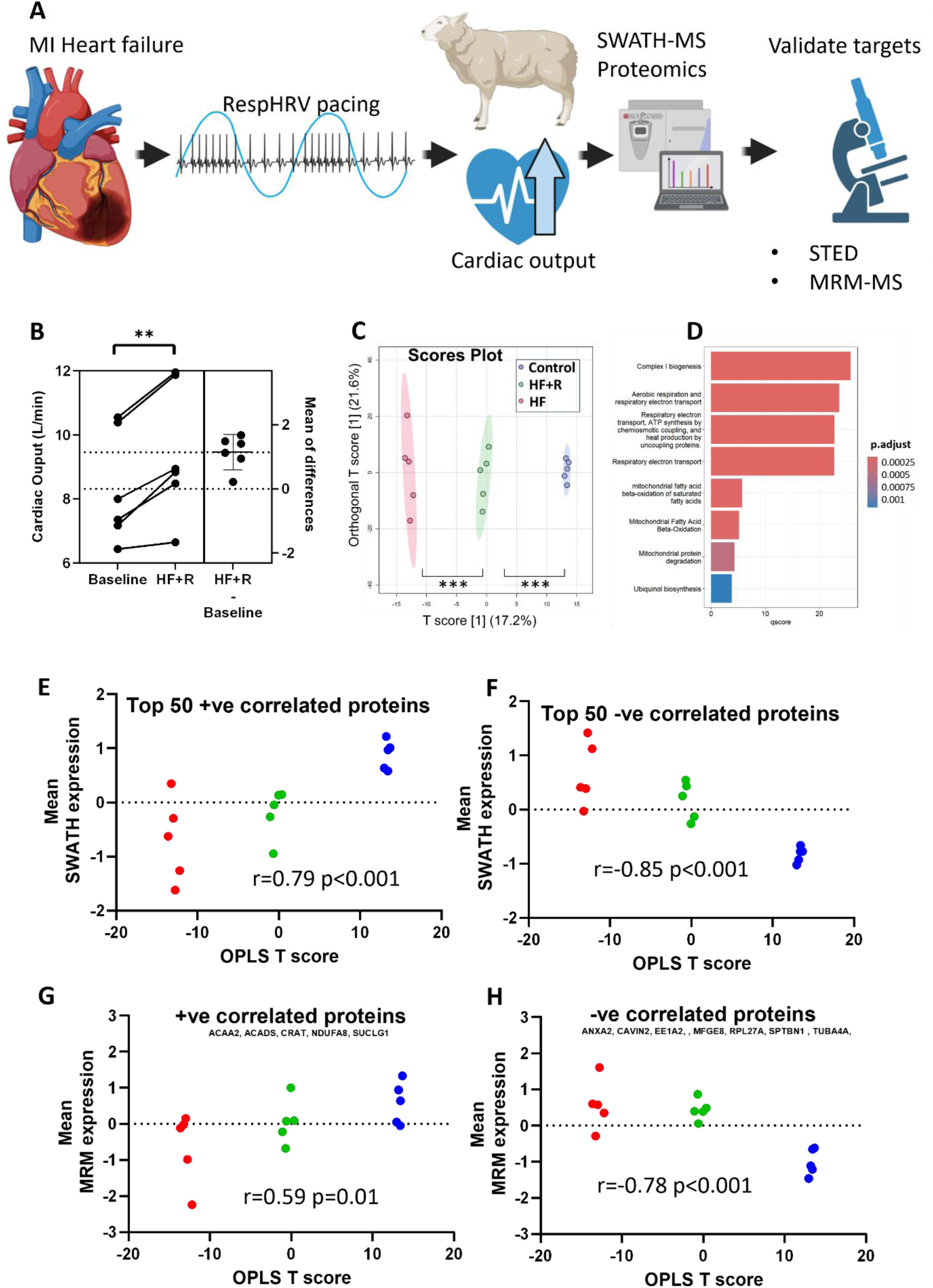
Proteomics analysis of the failing sheep heart demonstrates restoration of the expression of mitochondrial proteins and reduced expression of largely cytoskeletal proteins after 2 weeks of RespHRV pacing. **A** Graphical representation of the experiment approach that used SWATH-MS to identify key target proteins that changed expression after RespHRV pacing, and these targets were then validated with resolution microscopy and MRM-MS. **B** Paired T-test showed a significant (***p<0.001) change in cardiac output after 2 weeks of RespHRV pacing in heart failure sheep (HF+F). **C** OPLS analysis of the SWATH mass spectrometry data showed a clear separation between the groups; control (C), heart failure (HF), and HF+R. A one-way ANOVA of OPLS T score followed by Bonferroni corrected t-test showed a significant (***p<0.001) difference between the groups . **D** A Pathway Enrichment Analysis of the top 50 positively correlated proteins (down in HF) found a significant (p<0.001***) over-representation of mitochondrial proteins. **E** The mean SWATH expression of the top 50 positively correlated proteins demonstrated a significant Pearson’s correlation with the OPLS T score (r=0.74 p<0.001***). **F** The top 50 negatively correlated proteins (up in heart failure) were dominated by cytoskeletal proteins with their mean SWATH expression showing a significant Pearson’s correlation with the OPLS T score (r=-0.90 p<0.001***). MRM analysis of selected protein targets confirmed the SWATH results. **G** The mean MRM expression of the five validated positively correlated (mitochondrial) proteins showed a significant Pearson’s correlation with the SWATH OPLS T score (r= 0.59 p=0.008**). **H** The mean MRM expression of the eight negatively correlated (largely cytoskeletal) proteins showed a significant Pearson’s correlation with the OPLS T score (r=-0.75 p<0.001***). For panel B, n=6 HF+R sheep. For the proteomics n = 5 sheep for each experimental group (C, HF, HF+R).

### Tissue collection

The sheep were euthanized with an overdose of sodium pentobarbitone (0.5 ml/kg, IV, Provet NZ Pty Ltd., New Zealand). The heart was placed in cold PBS and rinsed twice. For proteomics, a portion of LV was frozen in liquid nitrogen and stored at -80°C. For immunohistochemistry, a portion of LV wall was placed in Tyrode’s solution (137 mM NaCl, 2.7 mM KCl, 1 mM CaCl_2_, 1 mM MgCl_2_, 0.2 mM Na_2_HPO_4_, 12 mM NaHCO_3_, 5.5 mM D-glucose) with 20 mM 2,3-butanedione monoxime (BDM) at room temperature for 10 minutes. Samples were then transferred to 1% paraformaldehyde (PFA) solution in PBS for one hour, at 4 degrees, before being moved through three solutions of increasing sucrose gradient (10%, 20%, 30%) over 48h. Once cryoprotected with sucrose, tissue samples were frozen in liquid nitrogen chilled methylbutane (Thermo Fisher Scientific, USA), and stored at -80°C.

### Proteomics

To each heart tissue sample 150 µl of 7M urea, 2M Thiourea, 5mM DTT, 0.1% SurfactAmps X-100 in 50 mM ammonium bicarbonate was added, and the samples probe sonicated for a total of 3 minutes. Disulphide bonds were reduced by incubation at 56°C for 20 minutes. Cysteines were alkylated by the addition of iodoacetamide (IAM) to 15 mM final concentration, and the samples were incubated in the dark at room temperature for 30 min, followed by the addition of cysteine to 15 mM final concentration to quench residual IAM. Total protein content was determined using the EZQ Protein Quantitation kit (Thermo Fisher Scientific, USA) as per the manufacturer’s instructions. An aliquot corresponding to 20 µg of total protein was taken from each sample and diluted 10-fold with 50 mM ammonium bicarbonate prior to addition of 0.5 µg sequencing grade modified porcine trypsin (Promega, Madison, WI, USA), and the samples incubated at 45°C for 1 hr in a chilled microwave (CEM, Matthews, NC, USA) using 15W power. Subsequently, an additional 0.5ug of trypsin was added to each sample followed by incubation overnight at 37°C in an incubator (Mini Incubator, Labnet, Woodbridge, NJ, USA). After digestion, samples were acidified to pH 3 via the addition of 50% formic acid, centrifuged for 3 min at 16,000g, and desalted using 10 mg Oasis HLB SPE cartridges (Waters, Milford, MA, USA) as per the manufacturer’s instructions. Purified peptides were eluted with 300 µl of 50% acetonitrile in 0.1% formic acid, and then concentrated to a final volume of 25 µl in a vacuum concentrator.

Spectral library sample analysis was carried out via data-dependent acquisition (DDA) mode: For the SWATH Ion Library generation step, a pool was made of a subsample of each of the above digested samples, and applied to an SCX MicroSpin column (The Nest Group, Inc.) according to manufacturer’s instructions and fractionated using 50 mM, 60 mM, 70 mM, 80 mM, 90 mM, 100 mM, 120 mM, 140 mM, 160 mM, 180 mM, 200 mM, and 500 mM NaCl. The collected fractions were desalted on 10 mg Oasis HLB SPE cartridges as above, and vacuum concentrated to 20 µl. The LC-MS/MS analysis of these fractions, along with one unfractionated sample, was performed on an Eksigent 425 nanoLC chromatography system (Sciex, Framingham MA, USA) connected to a TripleTOF 6600 Quadrupole-Time-of-Flight mass spectrometer (Sciex). Samples were injected onto a 0.3x 10mm trap column packed with Reprosil C18 media (Dr Maisch) and desalted for 5 minutes at 10ul/min before being separated on a 0.075 x 200 mm picofrit column (New Objective) packed in-house with Reprosil 3u C18-AQ media. The following gradient was applied at 300 nl/min using a NanoLC 400 UPLC system (Eksigent): 0 min 1% B; 2min 5% B; 47 min 35% B; 48 min 95% B; 50.5 min 95% B; 51 min 1% B; 60 min 1% B where A was 0.1% formic acid in water and B was 0.1% formic acid in acetonitrile. The picofrit spray was directed into a TripleTOF 6600 mass spectrometer (Sciex) operating in Information Dependent Acquisition (IDA) mode; scanning from 300-1600 m/z for 200 ms, followed by 35 ms MS/MS scans on the 40 most abundant multiply-charged peptides (m/z 80-1600) for a total cycle time of ∼1.65 seconds. The mass spectrometer and HPLC system were under the control of the Analyst TF 1.7 software package (Sciex).

#### Spectral library generation

The resulting data was searched against a database comprising the sheep proteome (downloaded from Uniprot.org) appended with a set of common contaminant sequences (23,265 entries in total) using Fragpipe. Search parameters were as follows: Iodoacetamide was set as global cysteine alkylation modification. Trypsin was selected as the digestion enzyme with up to 2 mis-cleavages. The precursor and product ion tolerances were set as 20 ppm. The Percolator tool was used to validate PSM results. ProteinProphet was used to validate the Protein level results. FDR filter set at 1% was applied to the search result report. The resulting group file used MSBooster to summarize and predict the spectra retention time. The Spectral library was then exported to a TSV file for SWATH data quantification.

#### SWATH sample acquisition and data analysis

SWATH analysis was then conducted for each sample using an acquisition method comprising 100 variable width isolation windows (with 1 Da overlap) covering a precursor mass range of 350–1160 m/z. The accumulation time was 50ms for the initial TOF-MS scan (m/z 350-1600), and 15ms for each SWATH MS/MS scan (m/z 140-1600), giving a total cycle time of ∼1.6s, employing the same LC conditions as described above, on the same mass spectrometer. DIA-NN was used for the SWATH data quantification^15^. The constructed spectral library was converted to the DIA-NN compatible library via Fragpipe^16^. The match-between-runs method was selected to produce more consistent quantification results. Normalization at the injection level was disabled. The mass accuracy was automatically estimated using the first run in the experiment. Cross-run analysis was conducted to generate global q-values for protein and gene groups. Output was filtered at 1% FDR for precursors and protein/gene groups. A report spectral library was re-assembled from the first pass result. The mass accuracy and RT window were re-estimated using the runs in the experiment. The peaks were picked and integrated in double-pass mode of the Neural network classifier.

#### Targeted Proteomic Analysis (MRM-HR)

A list of m/z values with corresponding retention times for a set of 221 peptide sequences of interest was compiled and used as an “inclusion” list within an IDA method targeting the top 55 features in each cycle. The accumulation time was 150ms for the initial TOF-MS scan (m/z 300-1600), and 35ms for each MS/MS scan (m/z 140-1600), giving a total cycle time of ∼2.1s. The same LC hardware, solvent gradient, and mass spectrometer as described above were used for this analysis. The raw data were then imported to Skyline (daily 23.1.1.425) for the targeted precursor quantitation. Peak area was normalized by the beta actin protein in the target list. The proteomic data set can be accessed at https://www.ebi.ac.uk/pride/login using the Reviewer Token: PXD057595 and password: QrDflz2Svmwq.

### Immunohistochemistry, and STED microscopy

LV sections 10 µm thick were cut using a Cryostat (CM3050, Leica, Germany) and mounted onto poly-L-lysine coated glass coverslips (22 x 50 mm, no. 1.5, Menzel™, Thermo Fisher Scientific, USA), which were taped on to labelled 25 x 75 x 1 mm microscope glass slides and stored at -80°C. Tissue sections were permeabilised in 1% Triton x-100 in PBS for 15 min and then blocked in Image-iT® FX signal enhancer (Thermo Fisher Scientific, USA) for 60 min at room temperature (RT). Sections were then incubated with either of the following primary antibody pairs; rabbit anti-Tomm20 (1:100, SC-11415, Santa Cruz Biotechnology) and mouse anti-RyR (1:100, MA3-916, Thermo Fisher Scientific, USA), or rabbit anti-collagen VI (1:100, SC-11415, Abcam, UK) and mouse anti-RyR (MA3-916, Thermo Fisher Scientific, USA) diluted in incubation solution (1%BSA, 0.05% Triton x-100, 0.05% NaN3 in PBS) overnight at 4°C. The following day, tissue sections were washed in PBS and then incubated in a solution containing the secondary antibody goat anti-rabbit Alexa Flour 594 (Thermo Fisher Scientific, USA) and goat anti-mouse Abberior STAR RED (Abberior, Germany) diluted 1:200 and incubated for 2 hours at RT. Sections were washed in PBS and mounted in ProLong Gold mounting medium (Thermo Fisher Scientific, USA) on a glass slide and poly-L-lysine coated #1.5 coverslips.

Two-dimensional STED images were obtained using an Olympus IX83 Abberior Facility Line STED microscope (Abberior Instruments, Germany) at the Biomedical Imaging Research Unit, University of Auckland, equipped with a 63x 1.4 NA oil immersion objective. Before each imaging session, the STED microscope was allowed to warm up, and routine alignment checks were conducted using Autoalignment Slide (Abberior Instruments, Germany). All images were collected from longitudinally orientated cardiomyocytes with a pixel size of 10 x 10 nm. For imaging of Alexa Fluor 594 the excitation laser was 561 nm, pinhole 1 Airy unit, STED laser 775 nm, detector 588-698 nm, line accumulation of 3, and STED gating of 750 ps (+8 ns). For the Abberior Star Red the excitation laser was 640 nm, STED laser 775 nm, detector 650-755 nm, line accumulation of 3, and STED gating of 750 ps (+8 ns).

### Image analysis

STED images were deconvolved with Huygens Essential version 24.04 (Scientific Volume Imaging, The Netherlands, http://svi.nl). The angle of Tomm20 labelling was calculated using Fiji/ImageJ directionality plugin. Tomm20 labelling area was calculated using a custom Fiji/ImageJ macro that thresholded the images with the Isodata algorithm and measured the area of the resulting mask. Myocyte widths were calculated from confocal images of collagen VI labelling in longitudinally orientated cells using Fiji/ImageJ. The angle of collagen VI/T-tubules labelling was calculated using Fiji/ImageJ directionality plugin. RyR2 cluster analysis utilised custom code written in Python utilising the scipy ndimages library^17^. First, T-tubule and RyR2 images were segmented into binary masks using the pixel classification feature of the machine learning software Ilastik version 1.4. The T-tubular mask was used to segment RyR2 labelling into junctional and corbular (non-junctional) RyR2 clusters. From these masks, RyR2 tetramers per cluster and RyR2 cluster density per cell area were measured. A filter was applied to the measured sizes where all cluster areas below 1 RyR2 were excluded from the analysis. In addition, the number of cardiac junctions per cellular area and the average junctional area were measured from a mask created of the overlap between T-tubule and RyR2 masks. T-tubule and RyR2 images were analyzed for regularity in the frequency domain for T-power and R-power, respectively^18^. T-tubule diameter was calculated as previously described^19^. This involved using Fiji/ImageJ and extracting a line profile perpendicular to the T-tubule axis and then fitting a Gaussian function to the data and calculating the Full Width Half Max (FWHM) as a measure of diameter.

### Bioinformatics and statistics

Statistical analyses were carried out using GraphPad Prism (Version:9.3.1) unless otherwise stated. A p<0.05 is considered significant. Change of cardiac output after RespHRV pacing was tested with a paired T-test. The MetaboAnalystR package (Version 5.0)^20^ was used for the Orthogonal Partial Least Squares (OPLS) discriminant analysis on the proteomics data and the visualization of results. Peak intensities were normalized by the median value of each sample. The log transformation, feature level mean centering, and rescaling were applied to the normalized data. ANOVA followed by Bonferroni-corrected LSD was used to test for the difference between the groups. The top 50 positive and top 50 negative proteins correlated with OPLS T-score were then selected for pathway enrichment, respectively. The enrichment analysis for gene ontology (GO) biological process and cellular component was performed using the clusterprofiler R package^21^. The Benjamini–Hochberg correction for multiple testing using a false discovery rate (FDR) of 5% was applied to determine significant functional categories and pathways. Correlation analysis was used to test for association between the mean expression of the top 50 positively correlated proteins and the mean expression of the top 50 negatively correlated proteins and the SWATH OPLS T score. Similarly, correlation analysis was used to test for association between the mean MRM expression of the confirmed positively and negatively correlated proteins, respectively, with the SWATH OPLS T score. Correlation analysis using an FDR of 5% was used to determine which MRM targets were significantly associated with the SWATH OPLS T score. For image analysis, a linear mixed model was used with a two-level hierarchy [level 1: group (wildtype vs. Col6a1-/-) and level 2: random factor animal (myocytes per animal)] followed by pairwise comparison and Bonferroni correction was used to test for difference between groups in IBM SPSS Statistics (Version: 28.0.1.1)

## Results

### Protein expression changes

Our study involved RespHRV-pacing in HF sheep for 2 weeks (HF+R) and using proteomics to compare this group to LV samples from HF sheep without pacing (HF) and healthy controls. Findings were validated by further proteomic analysis and super-resolution microscopy; the experimental approach is outlined in Figure 1A. After 2 weeks of RespHRV pacing, there was a 1.1 ± 0.2 L/min increase in cardiac output (Figure 1B) in the HF+R group. The SWATH-MS proteomics analysis identified 963 unique proteins in the LV of the sheep heart across the three groups, heart failure (HF), heart failure with RespHRV pacing (HF+R) and controls. These data were analysed by Orthogonal Partial Least Squares (OPLS) discriminant analysis, a data reduction methodology similar to Principal Components Analysis, but rather than maximising the variance explained, it seeks to maximise separation between the groups. A two-dimensional plot of the OPLS T score and orthogonal T score demonstrates a clear separation of the three groups (Figure 1C). The greatest separation was between the control and HF group, with HF+R group in an intermediate position. There was significant difference in between the three animal groups for OPLS T score, as presented in the plot in Figure 1C.

The top 50 proteins positively correlated with the OPLS T score or, in other words, reduced in HF and were partially restored in HF+R, with each protein having a correlation of greater than 0.55 (Table 1). Of these proteins, 42 /50 were mitochondrial in origin. A pathway Enrichment Analysis confirmed a significant over-representation of mitochondrial proteins (Figure 1D; p<0.001). The mean SWATH expression of these proteins showed a significant positive correlation with the OPLS T score (Figure 1E; p<0.001). Conversely, the top 50 proteins negatively correlated with the OPLS T score, or in other words, increased in HF and reduced in HF+R (Table 2), had correlations of greater than 0.62 and were dominated by cytoskeletal proteins, particularly those associated with microtubules. The mean SWATH expression of these proteins showed a significant negative correlation with the OPLS T score (Figure 1F; p< 0.001). Overall, these results indicate that LV structural proteins are up-regulated in HF and are reduced by RespHRV.

**Table 1.**
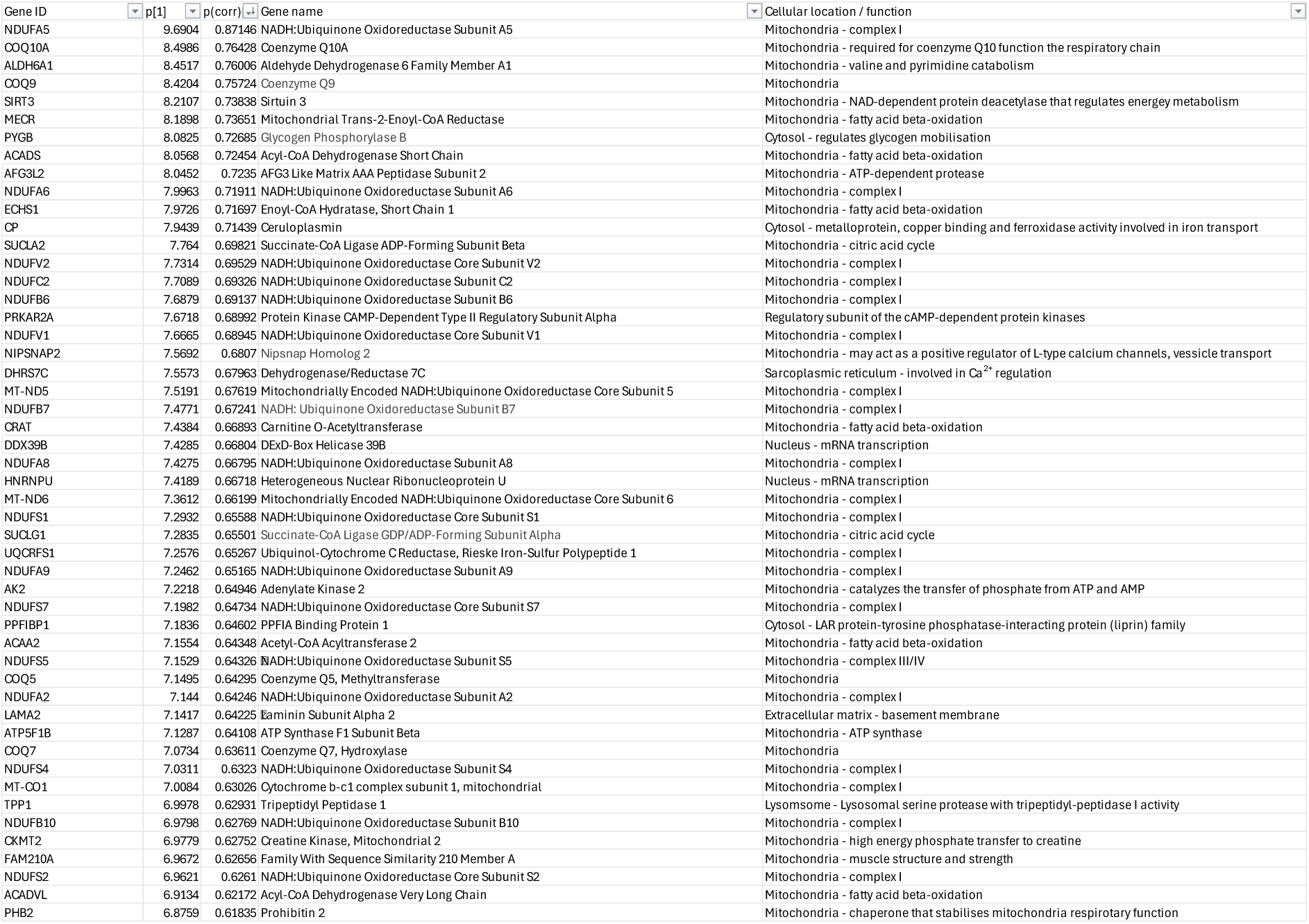
Proteins positively correlated with the SWATH OPLS-T score. Highest expression in control, intermediate expression in HF+R, and lowest expression in HF.

**Table 2.**
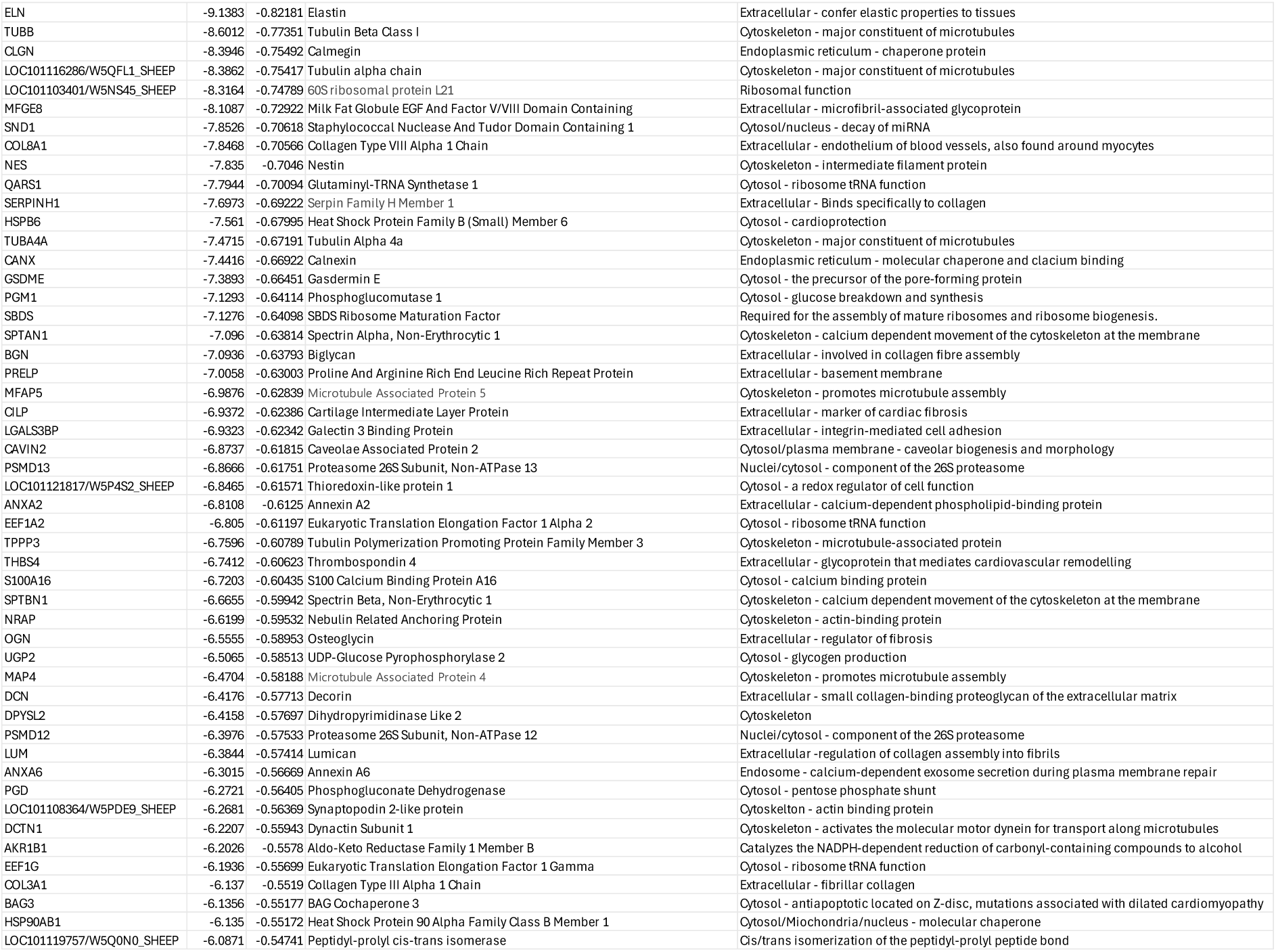
Proteins negatively correlated with the SWATH OPLS-T score. Highest expression in HF, intermediate expression in HF+R, and lowest expression in control.

The findings of the SWATH analysis were validated with the proteomic method of Multiple Reaction Monitoring (MRM). MRM works by targeting precursor ions of the proteins of interest and then selectively fragmenting the selected ions into product ions for further mass analysis. The technique has high specificity, accuracy, and reproducibility, providing a gold standard for protein quantification^22^. In total, 14 SWATH targets that included both mitochondrial and cellular cytoskeleton proteins were selected for further analysis by MRM. Of these targets, 12 were found to be significantly correlated with the SWATH OPLS T score (Table 3). The positively correlated proteins were all of mitochondrial origin; Acyl-CoA Dehydrogenase Short Chain (ACADS), Acetyl-CoA Acyltransferase 2 (ACAA2), Succinate-CoA Ligase GDP/ADP-Forming Subunit Alpha (SUCLG1), Carnitine O-Acetyltransferase (CRAT), and NADH: Ubiquinone Oxidoreductase Subunit A8 (NDUFA8). Their mean MRM expression, similar to the SWATH expression, was positively correlated with the SWATH OPLS T score (Figure 1G). The negatively correlated targets included cytoskeletal proteins; Spectrin Beta, Non-Erythrocytic 1 (SPTBN1), Tubulin Alpha 4a (TUBA4A), plasma membrane/Ca^2+^ signalling proteins: Caveolae Associated Protein 2 (CAVIN2), Annexin A2 (ANXA2), proteins involved in transcription; Eukaryotic Translation Elongation Factor 1 Alpha 2 (EEF1A2), Ribosomal Protein L27a (RPL27A) and extracellular matrix signalling; Milk Fat Globule EGF And Factor V/VIII Domain Containing (MFGE8). Their mean MRM expression, similar to the SWATH expression, was negatively correlated with the SWATH OPLS T score (Figure 1H).

**Table 3.**
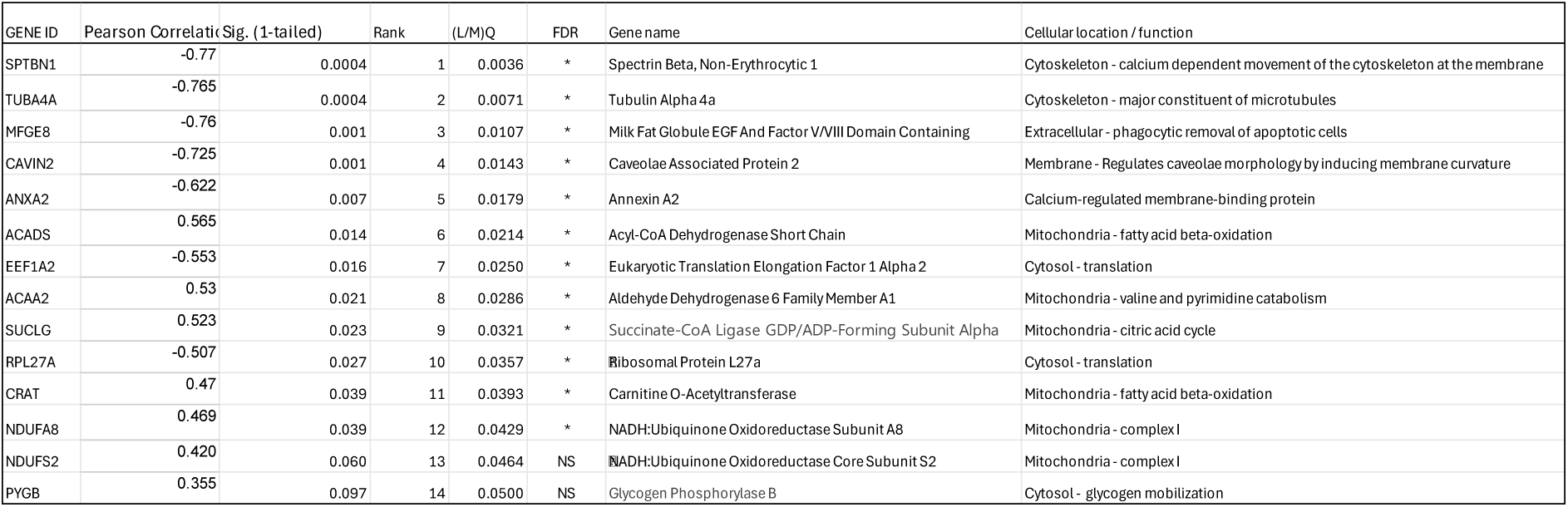
MRM proteomics of selected targets. FDR = false discovery rate set at 5%. * = significant after FDR correction. NS = non-significant.

| GENE ID | Pearson Correlation | Sig. (1-tailed) | Rank | (L/M/Q) | FDR | Gene name | Cellular location / function |
| --- | --- | --- | --- | --- | --- | --- | --- |
| SPTBN1 | -0.77 | 0.0004 | 1 | 0.0036 | * | Spectrin Beta, Non-Erythrocytic 1 | Cytoskeleton - calcium dependent movement of the cytoskeleton at the membrane |
| TUBA4A | -0.765 | 0.0004 | 2 | 0.0071 | * | Tubulin Alpha 4a | Cytoskeleton - major constituent of microtubules |
| MFGE8 | -0.76 | 0.001 | 3 | 0.0107 | * | Milk Fat Globule EGF And Factor V/VIII Domain Containing | Extracellular - phagocytic removal of apoptotic cells |
| CAVIN2 | -0.725 | 0.001 | 4 | 0.0143 | * | Caveolae Associated Protein 2 | Membrane - Regulates caveolae morphology by inducing membrane curvature |
| ANXA2 | -0.622 | 0.007 | 5 | 0.0179 | * | Annexin A2 | Calcium-regulated membrane-binding protein |
| ACADS | 0.565 | 0.014 | 6 | 0.0214 | * | Acyl-CoA Dehydrogenase Short Chain | Mitochondria - fatty acid beta-oxidation |
| EEF1A2 | -0.553 | 0.016 | 7 | 0.0250 | * | Eukaryotic Translation Elongation Factor 1 Alpha 2 | Cytosol - translation |
| ACAA2 | 0.53 | 0.021 | 8 | 0.0286 | * | Aldehyde Dehydrogenase 6 Family Member A1 | Mitochondria - valine and pyrimidine catabolism |
| SUCLG | 0.523 | 0.023 | 9 | 0.0321 | * | Succinate-CoA Ligase GDP/ADP-Forming Subunit Alpha | Mitochondria - citric acid cycle |
| RPL27A | -0.507 | 0.027 | 10 | 0.0357 | * | Ribosomal Protein L27a | Cytosol - translation |
| CRAT | 0.47 | 0.039 | 11 | 0.0393 | * | Carnitine O-Acetyltransferase | Mitochondria - fatty acid beta-oxidation |
| NDUFA8 | 0.469 | 0.039 | 12 | 0.0429 | * | NADH:Ubiquinone Oxidoreductase Subunit A8 | Mitochondria - complex I |
| NDUFS2 | 0.420 | 0.060 | 13 | 0.0464 | NS | NADH:Ubiquinone Oxidoreductase Core Subunit S2 | Mitochondria - complex I |
| PYGB | 0.355 | 0.097 | 14 | 0.0500 | NS | Glycogen Phosphorylase B | Cytosol - glycogen mobilization |

### Structure and position of mitochondria

Super-resolution imaging using STED microscopy, with 30 nm resolution, was used to assess the structure of the mitochondria across the three experimental groups using an antibody against the Translocase of Outer Mitochondrial Membrane 20 (TOMM20) and the cardiac calcium channel the Ryanodine Receptor-2 (RyR2; Figure 2). This analysis revealed a column-like organisation of the mitochondria in the control hearts that was disrupted in HF and restored in HF+R (Figure 2A-C). This structural feature was analysed using the ImageJ directionality plugin that determines the orientation of structures within an image (Figure 2G). This analysis revealed a significant loss of the 90° signal fraction in HF that was restored in the HF+R paced group (Figure 2H; p=0.051). Moreover, area fraction analysis of TOMM20 labelling showed a significant increase in mitochondria in the HF+R group compared to HF (Figure 2I). Interestingly, in HF, there was a high proportion of cells (12/25) that showed a concentration of TOMM20 labelling near the Z-line (Figure 2E). In HF+R, similar cells were seen at a lower frequency (7/25), and typically, there were prominent regions within these cells with strong labelling between the Z lines, (Figure 2F). This feature was only observed in one myocyte in the control hearts (1/25; Figure 2D).

**Figure 2.**
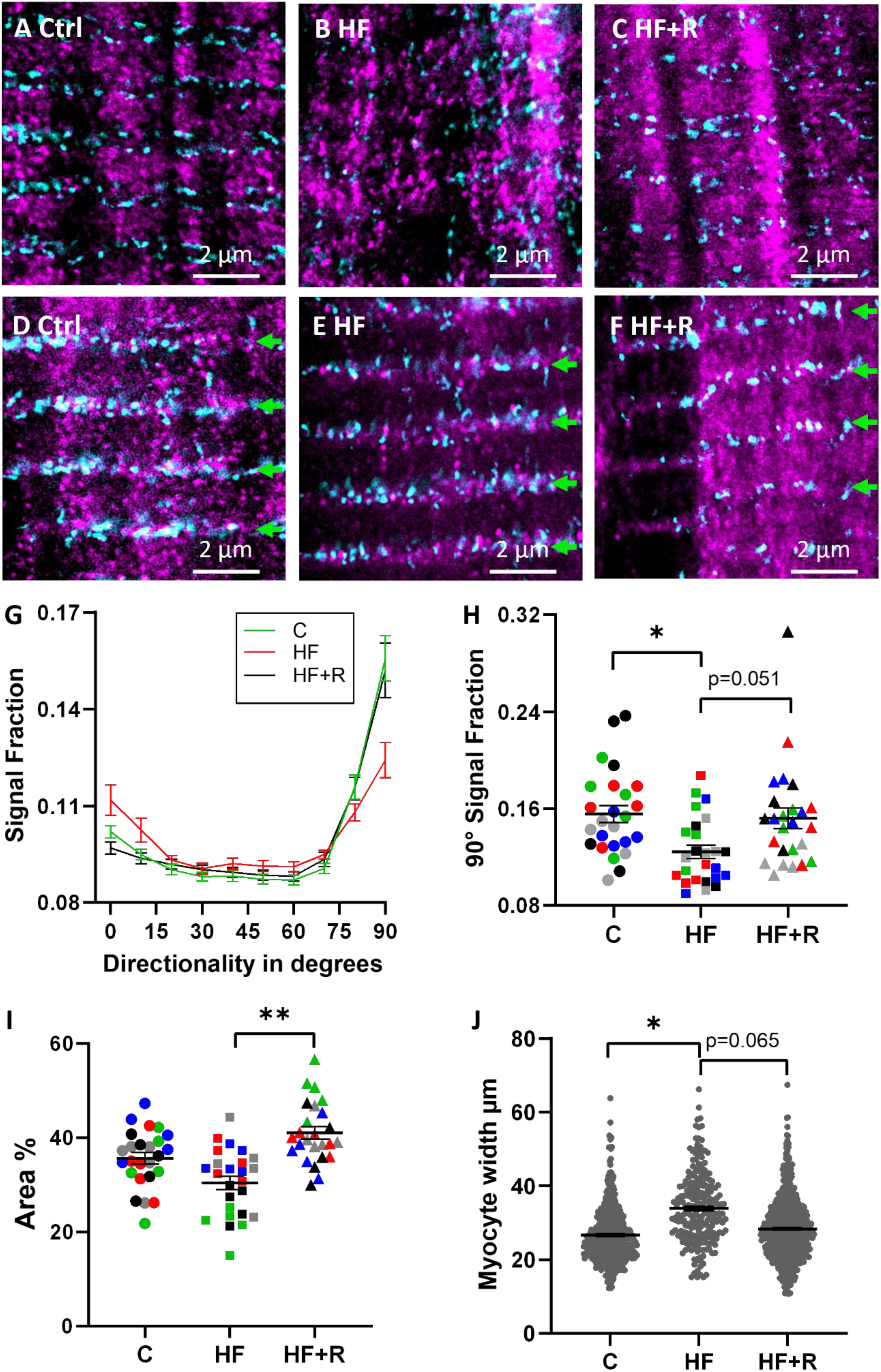
Super-resolution imaging of the failing sheep heart demonstrates restoration of the mitochondrial structure after 2 weeks of RespHRV pacing. STED images of longitudinally orientated myocytes labelled for TOMM20 (magenta) and RyR2 (cyan). **A** control (C). **B** heart failure without pacing (HF). **C** heart failure after RespHRV pacing (HF+R). Some cells demonstrated a concentration of TOMM20 at the Z-line, sample images are shown. **D** C. **E** HF. **F** HF+R (G). **G** Directionality analysis of TOMM20 labelling identified a dominant (90°) vertical alignment of the mitochondria. **H** The vertical alignment was lost in HF (*p=0.03) and restored in HF+R (p=0.051). **I** Area analysis of TOMM20 labelling showed an increase in mitochondria after RespHRV pacing (p=0.005). **G** The width of the myocytes was significantly wider in HF (*p=0.012) and treneded to smaller width in HF+R (p=0.065). Error bars are SEM of the mean. For panels H-I, each data point is the mean value for each cell analysed with symbols of the same colour from the same animal. For panels H-I, statistics used a mixed model analysis with two-level hierarchy (level 1: group (C, HF, HF+R), level 2: random factor animal (5 myocytes per animal). For panels L, statistics used a mixed model analysis with two-level hierarchy (level 1: group (C, HF, HF+R), level 2: random factor animal (16-152 myocytes per animal). For panels H-L comparisons were planned and used the LSD test, as the changes in mitochondria were predicted from the proteomics data and changes in myocyte hypertrophy were previously documented after 4 weeks RSA pacing^3^.

### T-tubule / Myocyte structure

Our previous work using confocal microscopy demonstrated that T-tubule structure was more intact and regular in HF sheep after 4 weeks of RespHRV pacing compared to HF sheep with 4 weeks of monotonic pacing^3^. Moreover, there was reduced myocyte hypertrophy in HF sheep after RespHRV pacing^3^. To explore the role of T-tubule reverse remodelling in response to 2-weeks of RespHRV pacing, higher resolution STED microscopy was used to image heart tissue labelled for collagen VI (T-tubules) and RyR2 (Figure 3A-F). STED images of HF tissue showed a noticeable loss of regular T-tubule structure that appeared to be mitigated in the HF+R group. Directionality analysis was used to assess the dominant direction of the T-tubules to quantify this apparent structural change (Figure 3A). This analysis demonstrated a significant loss of transverse T-tubules and an increase in axial T-tubules in HF (Figure 3H and 3I, respectively). There was weak evidence that RespHRV pacing partially reversed this trend, with the RespHRV group having a mean closer to the control group but was not significantly different from either the HF or control groups. Frequency analysis of T-tubule labelling (T-power) and RyR labelling (R-power) showed no change in the regularity of either of these labels, (Figure 3D, E). Moreover, analysis of T-tubule widths showed no difference between the groups (Figure 3F). Myocyte hypertrophy was assessed by measuring cellular widths in confocal images of collagen VI (Figure 3J). This analysis identified increased cell widths in HF, but after RespHRV pacing, although cellular width was decreased, this was not significantly different (p=0.065). Moreover, the HF+R was not significantly wider than the control group.

**Figure 3.**
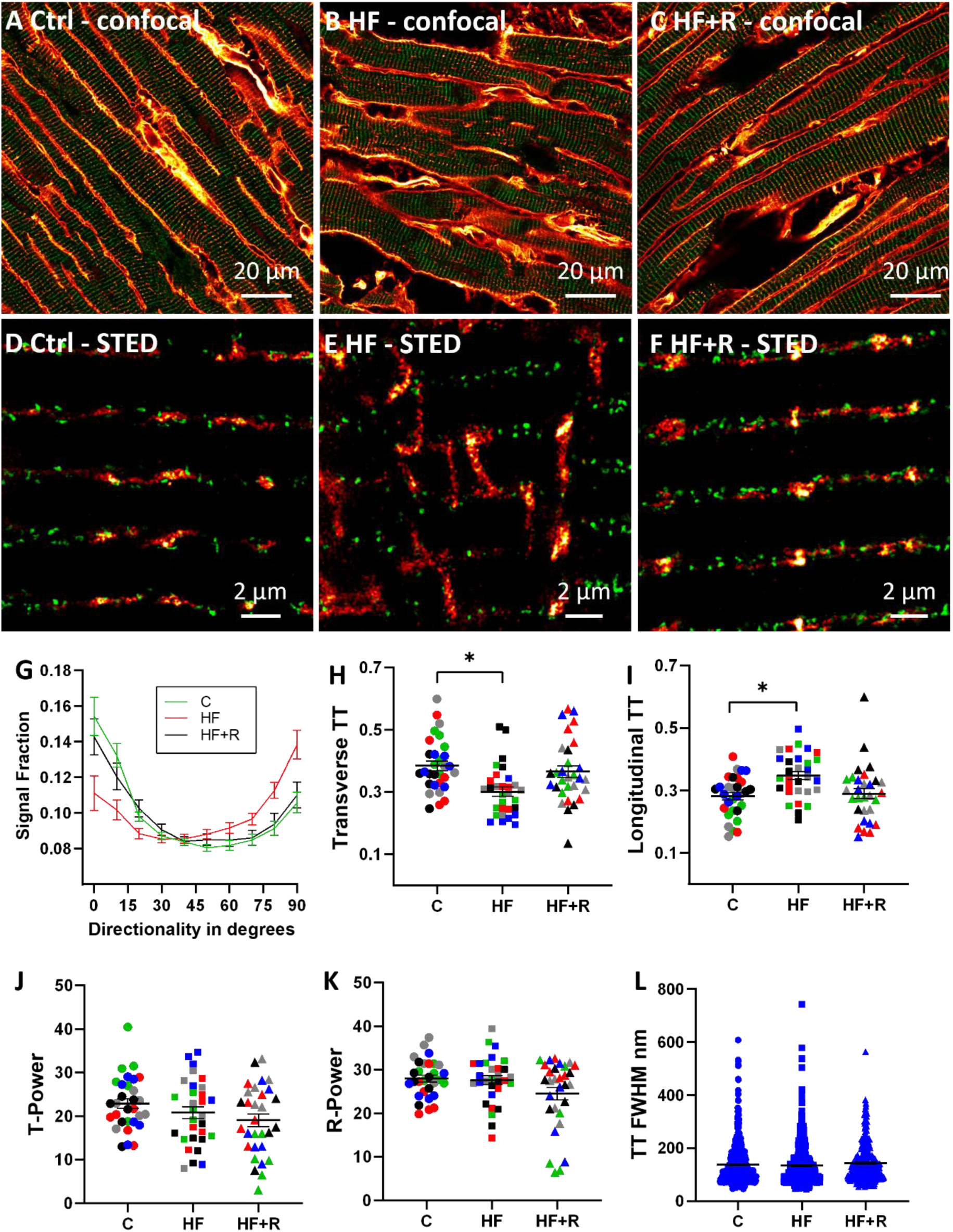
STED microscopy of Collagen VI and RyR2 in the failing sheep heart demonstrates a significant loss of T-tubule structure that is not reversed with 2 weeks of RespHRV pacing. **A-C** Confocal overview of longitudinally orientated myocytes labelled for collagen VI (red hot) and RyR2 (green) in control (panel A), heart failure without pacing (panel B) and heart failure after RSA pacing (panel C). **D-F** Examples of STED imaging of longitudinally orientated myocytes labelled for collagen VI (red hot) and RyR2 (green) in control (panel D), heart failure without pacing (panel E) and heart failure after RespHRV pacing (panel F). **G** The orientation of the t-tubules relative to the myocyte longitudinal axis was assessed with a directionality algorithm analysis of t-tubules labelled with collagen VI and showed both a transverse and axial component to the labelling. **H** The plot of the transverse (70-90°) component of the T-tubules, demonstrating a loss of transverse T-tubule in HF (*p=0.02). **I** The plot of the longitudinal (0-20°) component of the T-tubules showed increased axial tubules in HF (*p=0.02). **J** Frequency analysis of T-tubule labelling (collagen VI) showed no change between the groups. **K** Frequency analysis of the RyR2 labelling showed no change between the groups. **F** T-tubule width (FWHM) analysis of collagen VI labelling showed no change between the groups. Error bars are SEM of the mean. For panels H-K, each data point is the mean value for each cell analysed with symbols of the same colour from the same animal. For panels H-K, statistics used a mixed model analysis with two-level hierarchy (level 1: group (C, HF, HF+R), level 2: random factor animal (5 myocytes per animal). For panels L, statistics used a mixed model analysis with two-level hierarchy (level 1: group (C, HF, HF+R), level 2: random factor animal (43-152 T-tubles per animal). For panels H-L comparisons were planned and used the LSD test, as changes in T-tubule structure have been previously documented after 4 weeks of RSA pacing^3^.

### RyR2 cluster organisation

Previous research has shown that RyR2 clusters can become fragmented in HF^23–25^. To assess this feature, we examined the RyR2 and T-tubule STED images to assess the structure of both junctional and corbular RyR clusters (Figure 4). For this analysis, images of RyR2 and T-tubules were converted into a binary mask (Figure 4A-C). The RyR2 binary masks were used to calculate the number of RyR2 per cluster and the density of RyR2 clusters per cell, (Figure 5D, E). The T-tubule labelling was used to separate RyR2 clusters into junctional, or those that overlapped with T-tubule labelling, and corbular, those that were isolated from T-tubule labelling. This analysis showed there was no change in either RyR2 cluster size or density across the groups. However, in all groups, the junctional RyR2 cluster size was increased compared to corbular RyR2 clusters. There was also an increased number of corbular RyR2 clusters relative to junctional RyR2 clusters. Previous research has demonstrated a loss of cardiac junctions in HF^26^. To assess this feature, the co-localisation between the RyR2 and T-tubule masks was used to estimate the number and area of the junctions. This analysis demonstrated a significant increase in the number of cardiac junctions in HF compared to control hearts. However, there was no difference between the HF/RespHRV group with either HF or control hearts. The average area of the junctions was not different across the experimental groups.

**Figure 4.**
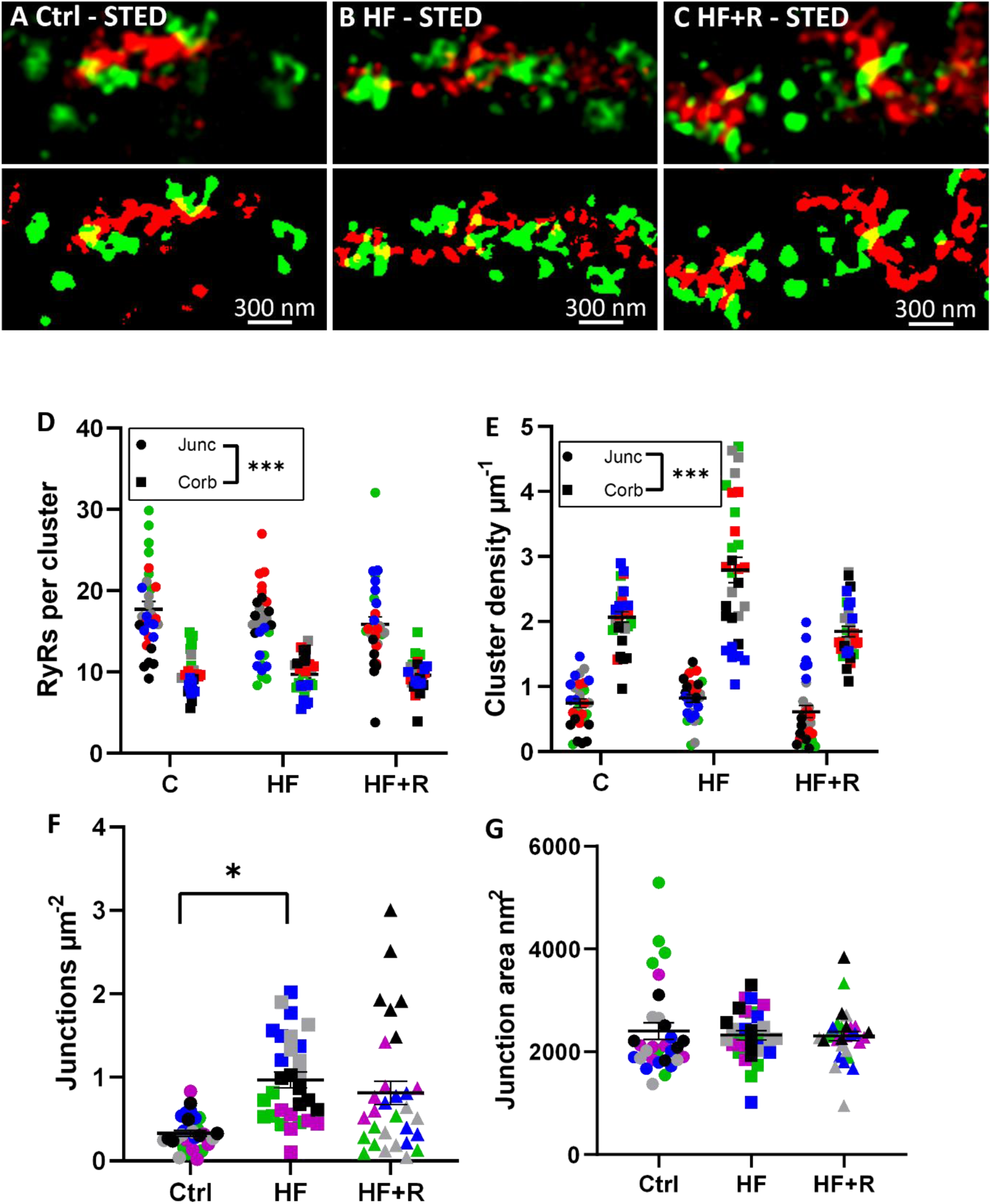
Analysis of RyR2 cluster organisation in the failing sheep heart demonstrates no significant improvement in the failing heart after 2 weeks of RespHRV pacing. **A-C** Zoomed-in representative images of RyR2 clusters (red), T-tubule collagen VI (green) and their colocalisation (yellow) for control (panel A), heart failure (panel B), and heart failure with RespHRV pacing (panel C). The top half of panels A-C shows exemplar STED imaging, and the bottom of panels A-C show the respective ilastik binary mask. The binary masks of the RyR2 clusters were separated into junctional RyR2 clusters (those that overlap with collagen VI) and corbular RyR2 clusters (those with no collagen VI overlap) for subsequent analysis. **D** The mean number of RyRs per cluster for control (C), heart failure (HF), and heart failure after RespHRV pacing (HF+R). **E** The mean RyR2 cluster density per cell area. **F** The mean junctional density. **G** The mean junction area (RyR2 and T-tubule collagen VI colocalisation). Error bars are SEM of the mean. Each data point is the mean value for each cell analysed with symbols of the same colour from the same animal. For this analysis, 5 hearts and 6 cells from each heart were analysed in each group (control, heart failure, and heart failure with RespHRV pacing). A linear mixed model with nesting of cells within the heart (random effect) was used to test for statistical significance between fixed effects; groups (control, heart failure and heart failure with RespHRV pacing) and cellular regions (junctional and corbular RyR2). Note for (F) and (G) that there is only one fixed effect, group (no cell region). For (D) and (E) the significance is p<0.001*** is the output from the mixed model for cell region. For (F) p=0.049* with Bonferroni correction.

**Figure 5.**
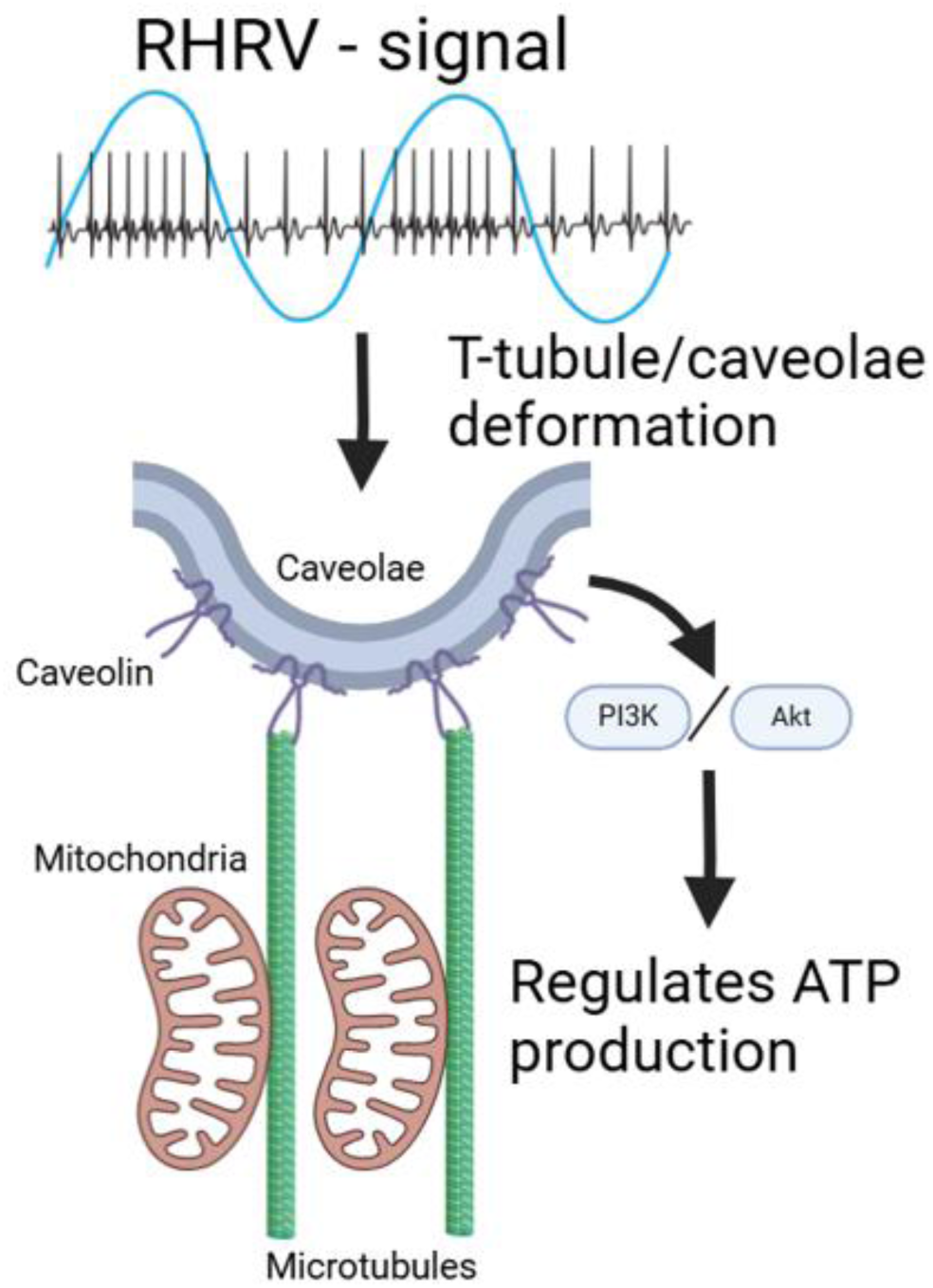
**Proposed mechanism. RespHRV signalling is used to coordinate contraction with mitochondrial ATP production through the myocyte caveolae/cytoskeleton network**.

## Discussion

Our study utilised multiple techniques to shed light on the mechanisms whereby RespHRV pacing exerts its beneficial actions. We show that RespHRV pacing significantly restores mitochondrial proteins involved in energy metabolism. We also show that several cytoskeletal proteins, particularly those of microtubule origin which are elevated in HF, were reduced after RespHRV pacing. Our objectives were investigated using SWATH-MS analysis, followed by targeted MRM-MS validation and finally by super-resolution STED microscopy. Taken together, our results suggest that RespHRV pacing results in beneficial actions on cardiac mitochondrial proteins which lead to reversal of cytoskeletal architecture in HF.

The first major finding of this study was that RespHRV pacing restored mitochondrial proteins involved in energy metabolism. This was demonstrated by the SWATH-MS analysis that identified a loss of mitochondrial proteins in HF that were restored partially with RespHRV pacing. The mitochondrial finding was validated by targeted MRM-MS that included proteins involved in complex I (NDUFA8), fatty acid beta-oxidation (ACADS, CRAT), citric acid cycle (SUCLG), and amino acid catabolism (ACAA2). Most of the mitochondrial proteins impacted were from complex I, which, when knocked down in the mouse heart, accelerates pressure overload-induced HF^27^. We followed the MRM-MS data with STED super-resolution of mitochondria which further supported this finding, showing that mitochondria structure was disorganised in HF and restored after RespHRV pacing, including increased volume of the mitochondria. Taken together, our data provide the first indication that the dramatic effects of RespHRV pacing may be mediated, in part, by the mitochondria. Dysfunctional mitochondrial respiration is a major contributor to CVD, with ATP levels reduced by ∼30% in HF^6^. In this context, our findings indicate that RespHRV may restore the energy balance in the failing heart. It is probable that by improving energetic efficiency, more energy is available to allow both repair and drive contraction; the latter explains the increased cardiac output after RespHRV pacing.

The second major finding of our proteomic study is that several cytoskeletal proteins, particularly those of microtubule origin (TUBB, Tubulin alpha chain, TUBA4A, MFAP5, TPP3, MAP4, DCTN1), were elevated in HF, but levels were reduced after RespHRV pacing. Interestingly, microtubule remodelling or increased levels of tubulin have been linked to cardiac T-tubule remodelling and Ca^2+^ handling defects in the dystrophic mouse^28^ and a mouse model of pressure overload HF^29^, which can be reversed in both models with tubulin depolymerisation. We suspect a similar mechanism is driving the T-tubule remodelling we have documented in HF sheep based on increases in microtubule proteins documented. Most relevant, the microtubule network is thought to regulate mitochondria metabolism through voltage-dependent anion channel (VDAC) located in the outer mitochondria membrane^30^. For example, in mouse cardiac myocytes, depolymerisation of either tubulin or actin cytoskeleton prevents L-type calcium channel agonist Bay K8644 from increasing mitochondrial membrane potential^31^. These data suggest an intriguing hypothesis that RespHRV regulates energy metabolism in the heart through direct mechanical linkage between the myocyte membrane and the mitochondria via the cytoskeleton (Figure 5). Further support for this hypothesis is provided by a study that showed mechanically stimulated Ca^2+^ release in rat myocytes is regulated by the spatial alignment of microtubules with the mitochondria^32^.

The proteomics data also identified SPTBN1 (βII spectrin) a member of the spectrin family of proteins that function to connect the cellular cytoskeleton to the membrane^33^. Recently, βII spectrin was shown to regulate cardiomyocyte mitochondria function with its conditional knockout triggering spontaneous contractile dysfunction and worsening cardiac function after ischaemia reperfusion^34^. Notably, spectrin has been found to be essential to mediate endothelial shear stress signalling via caveolae and the stretch-sensitive PIEZO1 channel^35^. Moreover, caveolae also regulate energy metabolism through interfacing with mitochondria^36^. Caveolae are present in the T-tubules at resting sarcomere length and become integrated (unfolded) into the T-tubules during both contraction and stretch^37^. This deformation of caveolae could provide a mechanism for the mechanical transmission of the RespHRV signal to myocyte mitochondria and the nucleus. Support for this mechanism comes from the proteomics identifying the caveolae protein CAVIN2, which has recently been shown to regulate PI3K-Akt signalling in cardiac myocytes^38^. Notably, PI3K-Akt signalling regulates metabolism, and its dysfunction leads to obesity and diabetes^39^. Furthermore, CAVIN2 has been shown to be localised to the T-tubules in mice with STED microscopy^40^.

The beneficial effects of RespHRV pacing could also involve other cell types in the heart. For example, ANAXA2, which was elevated in HF and reduced after RespHRV pacing in both the SWATH and MRM analysis, is a Ca^2+-^dependent phospholipid-binding protein that is thought to regulate actin cytoskeleton dynamics and is predominantly located within the capillaries, extracellular matrix, and endothelial cells of the coronary arteries in the human heart^41^. ANAXA2 expression is increased in human heart failure^42^ and serum levels are elevated in diabetic cardiomyopathy^43^. Although not generally associated with the myocytes, ANAXA2 is involved in skeletal muscle plasma membrane repair where it is part of the cap complex at the repair site^44^. Notably, ANAXA2 and caveolin-1 regulate pulmonary arterial smooth muscle proliferation^45^. Moreover, caveolin-1 expression in the heart is predominantly in the endothelial cells and smooth muscle cells^46^ but has been found in the cardiac myocyte membrane in mice^47^. Furthermore, caveolin-1 is involved in the mechanotransduction and remodelling of blood vessels^48^. A potential alternative mechanism of the response to RespHRV is through improved coronary blood flow that reduces hypoxic stress and allows the heart to function more efficiently. This hypothesis remains to be tested.

### Limitations and future directions

A limitation of this study is that we have shown an association between improved cardiac output and mitochondrial repair. The next step required is to experimentally test the hypothesis that RespHRV pacing improves energy metabolism. One approach would be to measure the energetics of the failing heart before and after RespHRV pacing and again after switching RespHRV pacing off. This could be done non-invasively by measurement of ATP and associated metabolites with the use of 31^P^ nuclear magnetic imaging^49^. Alternatively, equivalent measurements could be obtained from serial tissue analysis using endomyocardial biopsies as used clinically^50^. This would have the advantage of being able to assess mitochondrial respiration^51^ and measure cytoskeletal changes, providing a more in-depth test of the hypothesis.

Interestingly, we found weak evidence of reverse T-tubule and hypertrophic remodelling, suggesting that mitochondria effects are the dominant mechanism of the benefit provided by two weeks of RespHRV pacing. In comparison, in our previous study, 4 weeks of RespHRV pacing reduced both T-tubule remodelling and cardiac hypertrophy^3^, suggesting these t-tubule changes are downstream benefits to enhanced mitochondrial function. Surprisingly, cardiac junction density was increased in our model of HF, which is contrary to what has been observed by TEM in human end-stage heart failure^26^. A potential explanation is that increased junction density is a compensatory mechanism at an early stage in the heart failure trajectory to increase contractility in the heart. The lack of RyR2 cluster fragmentation in our model suggests early HF compared to other animal models of HF where RyR2 cluster fragmentation is a common finding^23–25^. Moreover frequency analysis commonly used to measure T-tubule remodelling failed to find differences in T-tubule structure between the animal groups, providing further support for an early HF phenotype. An interesting mitochondria feature observed that would be worthy of future investigation is the movement of TOMM20 towards the sarcoplasmic reticulum (RyR2) in the failing heart that was partially reversed with RespHRV pacing. TOMM20 regulates the importation of nuclear-encoded mitochondria proteins^52^ and this movement may facilitate protein uptake and indicate increased demand for protein importation that is required for repair of metabolically stressed mitochondria. The movement might also impact energy supply as it has been proposed that SR-mitochondria Ca^2+^ signalling regulates myocyte energy supply^53^.

## Conclusion

The results of this study using multiple techniques clearly show that RespHRV pacing increases the expression of mitochondrial proteins and that this change is associated with improved organisation and abundance of mitochondria, thus providing a plausible mechanism for how this novel form of pacing improves cardiac output in the failing heart. The decrease in cytoskeletal/caveolae proteins after pacing suggests that RespHRV is a signalling mechanism that coordinates the energy supply with the demands of contraction. There is extensive literature showing that the cytoskeleton is linked to the regulation of mitochondria^30–32,36^, and it is plausible that improved regulation of the mitochondria to the demand of the contraction through RespHRV pacing would produce energetic efficiencies. By improving energetic supply, energy can be directed to protein expression for cardiac work plus reparative processes. Future work will need to explore how the link between mitochondrial respiration and the cellular cytoskeleton is regulated by RespHRV pacing.

## References

1. La Rovere, M. T., Bigger, J. T., Marcus, F. I., Mortara, A. & Schwartz, P. J. Baroreflex sensitivity and heart-rate variability in prediction of total cardiac mortality after myocardial infarction. Lancet 351, 478–484 (1998).

2. Farmer, D. G. S., Dutschmann, M., Paton, J. F. R., Pickering, A. E. & McAllen, R. M. Brainstem sources of cardiac vagal tone and respiratory sinus arrhythmia. J Physiol 594, 7249–7265 (2016).

3. Shanks, J., Abukar, Y., Lever, N. A., Pachen, M., LeGrice, I. J., Crossman, D. J., Nogaret, A., Paton, J. F. R. & Ramchandra, R. Reverse re-modelling chronic heart failure by reinstating heart rate variability. Basic Res Cardiol 117, (2022).

4. O’Callaghan, E. L., Lataro, R. M., Roloff, E. L., Chauhan, A. S., Salgado, H. C., Duncan, E., Nogaret, A. & Paton, J. F. R. Enhancing respiratory sinus arrhythmia increases cardiac output in rats with left ventricular dysfunction. Journal of Physiology 598, 455–471 (2020).

5. Ben-Tal, A., Shamailov, S. S. & Paton, J. F. R. Central regulation of heart rate and the appearance of respiratory sinus arrhythmia: New insights from mathematical modeling. Math Biosci 255, 71–82 (2014).

6. Doenst, T., Nguyen, T. D. & Abel, E. D. Cardiac metabolism in heart failure: implications beyond ATP production. Circ Res 113, 709–724 (2013).

7. Levy D, G. R. S. D. K. W. C. W. Prognostic implications of echocardiographically determined left ventricular mass in the Framingham Heart Study. N Engl J Med 10, 1561– 1566 (1990).

8. Crossman, D. J., Young, A. A., Ruygrok, P. N., Nason, G. P., Baddelely, D., Soeller, C. & Cannell, M. B. t-tubule disease: Relationship between t-tubule organization and regional contractile performance in human dilated cardiomyopathy. J Mol Cell Cardiol 84, 170– 178 (2015).

9. Mellor, N. G., Pham, T., Tran, K., Loiselle, D. S., Ward, M., Taberner, A. J., Crossman, D. J. & Han, J. Disruption of transverse-tubular network reduces energy efficiency in cardiac muscle contraction. Acta Physiologica 231, e13545 (2021).

10. Han, J. C., Tran, K., Johnston, C. M., Nielsen, P. M. F., Barrett, C. J., Taberner, A. J. & Loiselle, D. S. Reduced mechanical efficiency in Left-Ventricular trabeculae of the spontaneously hypertensive rat. Physiol Rep 2, (2014).

11. Abukar, Y., Lever, N., Pachen, M., LeGrice, I. J. & Ramchandra, R. Impaired Baroreflex Function in an Ovine Model of Chronic Heart Failure Induced by Multiple Coronary Microembolizations. Front Physiol 10, 1420 (2019).

12. Shanks, J., Abukar, Y., Lever, N. A., Pachen, M., LeGrice, I. J., Crossman, D. J., Nogaret, A., Paton, J. F. R. & Ramchandra, R. Reverse re-modelling chronic heart failure by reinstating heart rate variability. Basic Res Cardiol 117, (2022).

13. O’Callaghan, E. L., Chauhan, A. S., Zhao, L., Lataro, R. M., Salgado, H. C., Nogaret, A. & Paton, J. F. R. Utility of a novel biofeedback device for within-breath modulation of heart rate in rats: A quantitative comparison of vagus nerve vs. right atrial pacing. Front Physiol 7, 1–13 (2016).

14. Nogaret, A., Zhao, L., Moraes, D. J. A. & Paton, J. F. R. Modulation of respiratory sinus arrhythmia in rats with central pattern generator hardware. J Neurosci Methods 212, 124– 132 (2013).

15. Demichev, V., Messner, C. B., Vernardis, S. I., Lilley, K. S. & Ralser, M. DIA-NN: neural networks and interference correction enable deep proteome coverage in high throughput. Nat Methods 17, 41–44 (2020).

16. Yu, F., Teo, G. C., Kong, A. T., Fröhlich, K., Li, G. X., Demichev, V. & Nesvizhskii, A. I. Analysis of DIA proteomics data using MSFragger-DIA and FragPipe computational platform. Nat Commun 14, (2023).

17. Hou, Y., Bai, J., Shen, X., de Langen, O., Li, A., Lal, S., dos Remedios, C. G., Baddeley, D., Ruygrok, P. N., Soeller, C. & Crossman, D. J. Nanoscale Organisation of Ryanodine Receptors and Junctophilin-2 in the Failing Human Heart. Front Physiol 12, 1649 (2021).

18. Moammer, H., Bai, J., Jones, T. L. M., Ward, M., Barrett, C. & Crossman, D. J. Pirfenidone increases transverse tubule length in the infarcted rat myocardium. Interface Focus 13, 20230047 (2023).

19. Crossman, D. J., Ruygrok, P. R., Soeller, C. & Cannell, M. B. Changes in the organization of excitation-contraction coupling structures in failing human heart. PLoS One 6, e17901 (2011).

20. Pang, Z., Zhou, G., Ewald, J., Chang, L., Hacariz, O., Basu, N. & Xia, J. Using MetaboAnalyst 5.0 for LC–HRMS spectra processing, multi-omics integration and covariate adjustment of global metabolomics data. Nat Protoc 17, 1735–1761 (2022).

21. Wu, T., Hu, E., Xu, S., Chen, M., Guo, P., Dai, Z., Feng, T., Zhou, L., Tang, W., Zhan, L., Fu, X., Liu, S., Bo, X. & Yu, G. clusterProfiler 4.0: A universal enrichment tool for interpreting omics data. Innovation 2, (2021).

22. Kulyyassov, A., Fresnais, M. & Longuespée, R. Targeted liquid chromatography-tandem mass spectrometry analysis of proteins: Basic principles, applications, and perspectives. Proteomics 21, e2100153 (2021).

23. Macquaide, N., Tuan, H. T. M., Hotta, J. I., Sempels, W., Lenaerts, I., Holemans, P., Hofkens, J., Jafri, M. S., Willems, R. & Sipido, K. R. Ryanodine receptor cluster fragmentation and redistribution in persistent atrial fibrillation enhance calcium release. Cardiovasc Res 108, 387–398 (2015).

24. Kolstad, T. R., van den Brink, J., Macquaide, N., Lunde, P. K., Frisk, M., Aronsen, J. M., Norden, E. S., Cataliotti, A., Sjaastad, I., Sejersted, O. M., Edwards, A. G., Lines, G. T. & Louch, W. E. Ryanodine receptor dispersion disrupts Ca 2+ release in failing cardiac myocytes. Elife 7, e39427 (2018).

25. Sheard, T. M. D., Hurley, M. E., Smith, A. J., Colyer, J., White, E. & Jayasinghe, I. Three-dimensional visualization of the cardiac ryanodine receptor clusters and the molecular-scale fraying of dyads. Philosophical Transactions of the Royal Society B: Biological Sciences 377, (2022).

26. Zhang, H.-B., Li, R.-C., Xu, M., Xu, S.-M., Lai, Y.-S., Wu, H.-D., Xie, X.-J., Gao, W., Ye, H., Zhang, Y.-Y., Meng, X. & Wang, S.-Q. Ultrastructural uncoupling between T-tubules and sarcoplasmic reticulum in human heart failure. Cardiovasc Res 98, 269–276 (2013).

27. Karamanlidis, G., Lee, C. F., Garcia-Menendez, L., Kolwicz, S. C., Suthammarak, W., Gong, G., Sedensky, M. M., Morgan, P. G., Wang, W. & Tian, R. Mitochondrial complex i deficiency increases protein acetylation and accelerates heart failure. Cell Metab 18, 239–250 (2013).

28. Prins, K. W., Asp, M. L., Zhang, H., Wang, W. & Metzger, J. M. PRE-CLINICAL RESEARCH Microtubule-Mediated Misregulation of Junctophilin-2 Underlies T-Tubule Disruptions and Calcium Mishandling in Mdx Mice VISUAL ABSTRACT. J Am Coll Cardiol Basic Trans Science vol. 1 (2016).

29. Zhang, C., Chen, B., Guo, A., Zhu, Y., Miller, J. D., Gao, S., Yuan, C., Kutschke, W., Zimmerman, K., Weiss, R. M., Wehrens, X. H. T., Hong, J., Johnson, F. L., Santana, L. F., Anderson, M. E. & Song, L.-S. Microtubule-mediated defects in junctophilin-2 trafficking contribute to myocyte T-tubule remodeling and Ca2+ handling dysfunction in heart failure. Circulation 129, 1742–1750 (2014).

30. Solomon, T., Rajendran, M., Rostovtseva, T. & Hool, L. How cytoskeletal proteins regulate mitochondrial energetics in cell physiology and diseases. Philosophical Transactions of the Royal Society B: Biological Sciences 377, (2022).

31. Viola, H. M., Johnstone, V. P. A., Szappanos, H. C., Richman, T. R., Tsoutsman, T., Filipovska, A., Semsarian, C., Seidman, J. G., Seidman, C. E. & Hool, L. C. The Role of the L-Type Ca 2þ Channel in Altered Metabolic Activity in a Murine Model of Hypertrophic Cardiomyopathy. J Am Coll Cardiol Basic Trans Sci 1, 61–72 (2016).

32. Miragoli, M., Sanchez-Alonso, J. L., Bhargava, A., Wright, P. T., Sikkel, M., Schobesberger, S., Diakonov, I., Novak, P., Castaldi, A., Cattaneo, P., Lyon, A. R., Lab, M. J. & Gorelik, J. Microtubule-Dependent Mitochondria Alignment Regulates Calcium Release in Response to Nanomechanical Stimulus in Heart Myocytes. Cell Rep 14, 140–151 (2016).

33. Liem, R. K. H. Cytoskeletal integrators: The spectrin superfamily. Cold Spring Harbor Perspectives in Biology vol. 8 Preprint at 10.1101/cshperspect.a018259 (2016).

34. Yang, R., Ruan, B., Wang, R., Zhang, X., Xing, P., Li, C., Zhang, Y., Chang, X., Song, H., Zhang, S., Zhao, H., Zhang, F., Yin, T., Qi, T., Yan, W., Zhang, F., Hu, G., Wang, S. & Tao, L. Cardiomyocyte βII spectrin plays a critical role in maintaining cardiac function by regulating mitochondrial respiratory function. Cardiovasc Res (2024) doi:10.1093/cvr/cvae116.

35. Mylvaganam, S., Plumb, J., Yusuf, B., Li, R., Lu, C. Y., Robinson, L. A., Freeman, S. A. & Grinstein, S. The spectrin cytoskeleton integrates endothelial mechanoresponses. Nat Cell Biol 24, 1226–1238 (2022).

36. Foster, C. R., Satomi, S., Kato, Y. & Patel, H. H. The caveolar-mitochondrial interface: Regulation of cellular metabolism in physiology and pathophysiology. Biochem Soc Trans 48, 165–177 (2020).

37. Rog-Zielinska, E. A., Scardigli, M., Peyronnet, R., Zgierski-Johnston, C. M., Greiner, J., Madl, J., O’Toole, E. T., Morphew, M., Hoenger, A., Sacconi, L. & Kohl, P. Beat-by-Beat Cardiomyocyte T-Tubule Deformation Drives Tubular Content Exchange. Circ Res 128, 203–215 (2021).

38. Maruyama, N., Ogata, T., Kasahara, T., Hamaoka, T., Higuchi, Y., Tsuji, Y., Tomita, S., Sakamoto, A., Nakanishi, N. & Matoba, S. Loss of Cavin-2 destabilizes PTEN and enhances Akt signaling pathway in cardiomyocytes. Cardiovasc Res (2024) doi:10.1093/cvr/cvae130.

39. Huang, X., Liu, G., Guo, J. & Su, Z. Q. The PI3K/AKT pathway in obesity and type 2 diabetes. Int J Biol Sci 14, 1483–1496 (2018).

40. Peper, J., Kownatzki-Danger, D., Weninger, G., Seibertz, F., Pronto, J. R. D., Sutanto, H., Pacheu-Grau, D., Hindmarsh, R., Brandenburg, S., Kohl, T., Hasenfuss, G., Gotthardt, M., Rog-Zielinska, E. A., Wollnik, B., Rehling, P., Urlaub, H., Wegener, J., Heijman, J., Voigt, N., Cyganek, L., Lenz, C. & Lehnart, S. E. Caveolin3 Stabilizes McT1-Mediated Lactate/Proton Transport in Cardiomyocytes. Circ Res 128, E102–E120 (2021).

41. Camors, E., Monceau, V. & Charlemagne, D. Annexins and Ca2+ handling in the heart. Cardiovasc Res 65, 793–802 (2005).

42. Benevolensky, D., Belikova, Y., Mohammadzadeh, R., Trouvé, P., Oise Marotte, F., Oise Russo-Marie, F., Samuel, J.-L. & Le Charlemagne, D. Expression and Localization of the Annexins II, V, and VI in Myocardium from Patients with End-Stage Heart Failure. Lab Invest 80, 123–133 (2000).

43. He, Q., Zhu, J., Yang, G., Liu, X., Li, L., Wang, Y., Xiong, X., Zheng, Y., Zheng, H. & Qu, H. Serum Annexin A2 concentrations are increased in patients with diabetic cardiomyopathy and are linked to cardiac dysfunctions. Diabetes Res Clin Pract 195, 110196 (2023).

44. Demonbreun, A. R., Quattrocelli, M., Barefield, D. Y., Allen, M. V., Swanson, K. E. & McNally, E. M. An actin-dependent annexin complex mediates plasma membrane repair in muscle. Journal of Cell Biology 213, 705–718 (2016).

45. Liao, L., Zheng, B., Yi, B., Liu, C., Chen, L., Zeng, Z. & Gao, J. Annexin A2-modulated proliferation of pulmonary arterial smooth muscle cells depends on caveolae and caveolin-1 in hepatopulmonary syndrome. Exp Cell Res 359, 266–274 (2017).

46. Ratajczak, P., Damy, T., Heymes, C., Oliviéro, P., Marotte, F., Robidel, E., Sercombe, R., Boczkowski, J., Rappaport, L. & Samuel, J. L. Caveolin-1 and -3 dissociations from caveolae to cytosol in the heart during aging and after myocardial infarction in rat. Cardiovasc Res 57, 358–369 (2003).

47. Cho, W. J., Chow, A. K., Schulz, R. & Daniel, E. E. Caveolin-1 exists and may function in cardiomyocytes. Can J Physiol Pharmacol 88, 73–76 (2010).

48. Yu, J., Bergaya, S., Murata, T., Alp, I. F., Bauer, M. P., Lin, M. I., Drab, M., Kurzchalia, T. V., Stan, R. V. & Sessa, W. C. Direct evidence for the role of caveolin-1 and caveolae in mechanotransduction and remodeling of blood vessels. Journal of Clinical Investigation 116, 1284–1291 (2006).

49. Abdurrachim, D. & Prompers, J. J. Evaluation of cardiac energetics by non-invasive 31P magnetic resonance spectroscopy. Biochimica et Biophysica Acta - Molecular Basis of Disease vol. 1864 1939–1948 Preprint at 10.1016/j.bbadis.2017.11.013 (2018).

50. Caves, P. K., Stinson, E. B., Graham, a F., Billingham, M. E., Grehl, T. M. & Shumway, N. E. Percutaneous transvenous endomyocardial biopsy. JAMA : the journal of the American Medical Association 225, 288–91 (1973).

51. Power, A. S. C., Pham, T., Loiselle, D. S., Crossman, D. H., Ward, M. L. & Hickey, A. J. Impaired ADP channeling to mitochondria and elevated reactive oxygen species in hypertensive hearts. Am J Physiol Heart Circ Physiol 310, H1649–H1657 (2016).

52. Zhao, F. & Zou, M.-H. Role of the Mitochondrial Protein Import Machinery and Protein Processing in Heart Disease. Front Cardiovasc Med 8, 749756 (2021).

53. De la Fuente, S. & Sheu, S. S. SR-mitochondria communication in adult cardiomyocytes: A close relationship where the Ca 2+ has a lot to say. Archives of Biochemistry and Biophysics vol. 663 259–268 Preprint at 10.1016/j.abb.2019.01.026 (2019).

54. Ben-Tal, A., Shamailov, S. S. & Paton, J. F. R. Evaluating the physiological significance of respiratory sinus arrhythmia: Looking beyond ventilation-perfusion efficiency. Journal of Physiology 590, 1989–2008 (2012).

